# The highly developed symbiotic system between the solar-powered nudibranch *Pteraeolidia semperi* and Symbiodiniacean algae

**DOI:** 10.1101/2023.04.30.538878

**Authors:** Hideaki Mizobata, Kenji Tomita, Ryo Yonezawa, Kentaro Hayashi, Shigeharu Kinoshita, Kazutoshi Yoshitake, Shuichi Asakawa

**Affiliations:** Department of Aquatic Bioscience, Graduate School of Agricultural and Life Sciences, the University of Tokyo; Technology Advancement Center, Graduate School of Agricultural and Life Sciences, the University of Tokyo

**Keywords:** symbiosis, endosymbiosis, Symbiodiniaceae, nudibranch, zooxanthellae, algae, coral

## Abstract

The intricate coexistence of Symbiodiniacean algae with a diverse range of marine invertebrates underpins the flourishing biodiversity observed within coral reef ecosystems. However, the phenomenon of coral bleaching—precipitated by the dissolution of symbiotic relationships with Symbiodiniaceae—poses a significant threat to these ecosystems, thereby necessitating an urgent investigation into the underlying symbiotic mechanisms. The symbiosis between nudibranchs and Symbiodiniaceae has been identified as an efficacious model for examining these mechanisms, yet a comprehensive understanding of their histological structures and cellular processes remains elusive. In this study, we concentrate on the nudibranch host *Pteraeolidia semperi*, renowned for its exceptional symbiotic capabilities, and we elucidate its sophisticated symbiotic architecture. Furthermore, we delineate the bleaching process within the nudibranch, uncovering the associated cellular processes. Collectively, these findings furnish invaluable insights into the intricate relationship between nudibranchs and Symbiodiniaceae, thereby contributing to our understanding of the symbiosis that sustains these critical marine ecosystems.

## Introduction

Dinoflagellates belonging to the family Symbiodiniaceae, colloquially referred to as “zooxanthellae,” engage in symbiotic relationships with an array of shallow-water marine invertebrates, including cnidarians,^1^ mollusks,^2^ and sponges.^3^ Typically, these host organisms furnish the dinoflagellates with requisite inorganic compounds,^4^ and the algae reciprocate by supplying photosynthetic derivatives, such as saccharides and lipids, to their hosts.^5^ Among these symbiotic associations, coral– Symbiodiniaceae interactions have garnered considerable scientific interest. Corals rely on algal symbionts for the majority of their metabolic requirements, which in turn facilitate the formation of coral reefs.^6^ These complex structures serve as indispensable habitats for marine life, fostering thriving ecosystems teeming with myriad species.^7^ Additionally, coral reefs contribute significantly to the economies of tropical and subtropical nations, primarily through the tourism industry.^8^ However, recent years have witnessed escalating incidents of coral bleaching and subsequent mortality^9^ due to the disintegration of the symbiotic relationship, with predictions suggesting that by the end of the 21st century, all coral reefs may be exposed to long-term bleaching risks.^10^ Such bleaching events have been demonstrated to drastically compromise coral reef biodiversity,^11^ rendering this an urgent environmental concern. To safeguard these diverse ecosystems and the economies they bolster, a thorough understanding of the molecular mechanisms governing coral–Symbiodiniaceae symbiosis is crucial. Consequently, the adoption of a comprehensive research approach that encompasses not only corals but also alternative host organisms as laboratory models has been underscored as a key strategy to address this pressing issue.^12^

Nudibranchs belonging to the Cladobranchia suborder exhibit an intracellular symbiotic relationship with Symbiodiniaceae,^13^ in contrast to other molluscan hosts that engage in extracellular symbiosis.^14,15^ It is posited that these nudibranchs acquire their algal symbionts via the consumption of prey cnidarians, rather than through direct acquisition from the water column.^16^ The striking resemblance between intracellular symbiosis in nudibranchs and corals underscores the potential value of employing nudibranchs as model organisms for investigating the mechanisms underlying coral– algal symbiosis.^17,18^ Nonetheless, to date, only 2 molecular studies^19,20^ have been conducted on this subject, leaving much to be uncovered about the symbiotic mechanism. Furthermore, the nudibranch *Berghia stephanieae*, which was employed in these investigations, is considered to possess a rudimentary symbiotic capacity.^18,21^ Although this species can incorporate dinoflagellates into its cells, it can maintain them for only a brief duration,^16^ and the essential material exchange system between partners, which is integral to invertebrate–zooxanthellae symbiosis, may not be fully functional. To effectively decipher the molecular mechanisms underpinning symbiosis, it is imperative to develop an experimental design rooted in histological findings—utilizing a sea slug model with advanced symbiotic capabilities encompassing both efficient algal uptake and material exchange processes— while addressing the current lack of comprehensive systematic histological knowledge in this domain.

The genus Pteraeolidia represents a rare example of nudibranchs with experimentally verified “symbiotic ability,” signifying the translocation of photosynthetic products from algae to the host organism.^22^ This genus is regarded as one of the most highly symbiotic nudibranchs, with records indicating survival for over 200 days without feeding due to their reliance on photosymbiosis.^23^ The presence of white individuals within this genus during certain seasons suggests a reduced quantity of dinoflagellates.^22,24^ This phenomenon resembles coral bleaching, and deciphering the bleaching mechanism in this species could provide insights into the process of coral bleaching. The limited histological studies on *Pteraeolidia* suggest that these nudibranchs possess intricately branched structures, called “fine tubules,” permeating their entire bodies, and that their cells contain zooxanthellae.^13,23,24^ In one electron microscopic analysis, Rudman reported images showing the presence of Symbiodiniaceae within the epithelial cells of the digestive gland.^24^ In a separate study, Kempf conducted another electron microscopic analysis, leading to the discovery of isolated “carrier” cells seemingly associated with the host digestive gland and the identification of Symbiodiniaceae and lipid droplets within these cells.^25^ This evidence suggests that zooxanthellae may symbiotically coexist within the carrier cells of *Pteraeolidia*, providing photosynthetic products to the host in the form of lipid droplets. However, the specific cellular processes underpinning this symbiosis remain unexplored. To gain a more profound understanding of the symbiotic mechanism, comprehensive and high-resolution image analyses using multifaceted observation techniques are warranted.

In this study, we present the most comprehensive and accurate observations to date on the structure of algal symbiosis in nudibranchs, utilizing the highly symbiotic nudibranch *Pteraeolidia semperi* as a model organism. Moreover, by examining the bleaching process of the nudibranch, we elucidate the histological processes involved in the symbiotic relationship between nudibranchs and Symbiodiniaceae. The insights gleaned from this research will significantly contribute to our understanding of the molecular mechanisms governing nudibranch–algal symbiosis.

## Materials and Methods

### Sample collection and bleeding

Specimens of *P. semperi* were collected from the intertidal zone in Miura City, Kanagawa Prefecture, Japan on May 12 and October 5, 2021. Following collection, the specimens were maintained in an artificial seawater tank containing 33 g/L of Instant Ocean Premium (NAPQO, Ltd.) and equipped with an external filtration system. The tank environment was kept under constant light conditions and maintained at a water temperature of 25°C. No feeding was provided for the duration of the study period.

### Comparison of algal retention among tissues

Individual sea slugs were anesthetized using a 10% magnesium sulfate solution, subsequently fixed in 10% seawater formalin, and then transferred and preserved in 100% glycerin. After a period of 3 months, the specimens were dissected into 9 distinct regions: rhinophore, tentacle, cerata, foot, epidermis, genital organs, radula and stomach, penis, and testis and ovary. The dissected tissues were reconstituted with phosphate-buffered saline (PBS). For tissue transparency, the CUBIC Trial Kit (FUJIFILM Wako Pure Chemical Corporation) was employed according to the manufacturer’s protocol. During the transparency process, nuclear staining was performed using DAPI Solution (FUJIFILM Wako Pure Chemical Corporation). Chlorophyll autofluorescence and DAPI fluorescence were observed in multiple layers using a fluorescence microscope (KEYENCE BZX-810). The optimal focus layer, characterized by the highest brightness, was selected using the microscope’s “best focus” function.

### Observation of algal retention before and after bleaching

Four nudibranch specimens were subjected to a fasting period of 50 days (2 bleached and 2 unbleached), after which the cerata from each individual were excised and fixed using a 4% paraformaldehyde (PFA) solution. The processes of tissue transparency, staining, and microscopic observation were carried out utilizing the same techniques as those previously described.

### Optimal microscope and TEM sectioning

Cerata, rhinophores, and tentacles from 0-day fasted, 15-day fasted (unbleached), and 50-day fasted (bleached) individuals were excised and fixed in an aldehyde fixing solution (2% PFA, 2% glutaraldehyde, 50 mM HEPES, 33 g/L Instant Ocean Premium). Subsequently, the samples were washed with distilled water and post-fixed with 1% osmium tetroxide (Nisshin EM) at 4°C for 1 hour. Following further washing and dehydration using an ascending ethanol series, the tissues were replaced with propylene oxide (Wako) and ultimately embedded in epoxy resin (Quetol 812, Nisshin EM). Resin blocks were sectioned using an ultramicrotome (Leica, Ultracut UCT). Semi-ultrathin sections (500-nm thickness) were stained with toluidine blue (Diagonal GmbH & Co. KG.) and observed under an optical microscope (KEYENCE BZX-810). Ultrathin sections (60-nm thickness) were stained with 4% uranyl acetate solution (Bio-Rad) and Reynolds’ lead citrate (Wako), followed by examination using a transmission electron microscope (JEOL JEM-1010). Transmission electron microscopy (TEM) images were overlapped and merged using Image Composite Editor (Microsoft). Scale bars were manually generated referencing the original images.

### SEM observation

For 0-day fasted and 15-day fasted (unbleached) individuals, the cerata were subjected to aldehyde fixation, osmium tetroxide fixation, and 100% ethanol substitution following the same procedure described earlier. Tissues were then fractured in liquid nitrogen and replaced with *t*-butyl alcohol (Wako). Once the freeze-drying process was completed, samples were affixed to an observation stage and coated with platinum-palladium using an ion sputter coater (Hitachi High-Tech E-1030). Finally, the samples were examined using a scanning electron microscope (Hitachi High-Tech, SEM-S4800).

## Results

### Symbiodiniaceae retention among different tissues

We dissected nudibranchs into 9 sections to identify tissues retaining algae and examined them for the presence of zooxanthellae. In this study, we detected Symbiodiniaceae by observing the autofluorescence of chlorophyll within the tissues. In superficial tissues such as the rhinophore, tentacle, epidermis, cerata, and foot, algae were consistently observed throughout the tissue (Fig. 1a, b). Conversely, no zooxanthellae were detected in internal tissues, including the radula, stomach, penis, genital organs, and the testis and ovary (Fig. 1c, d).

**Figure 1.**
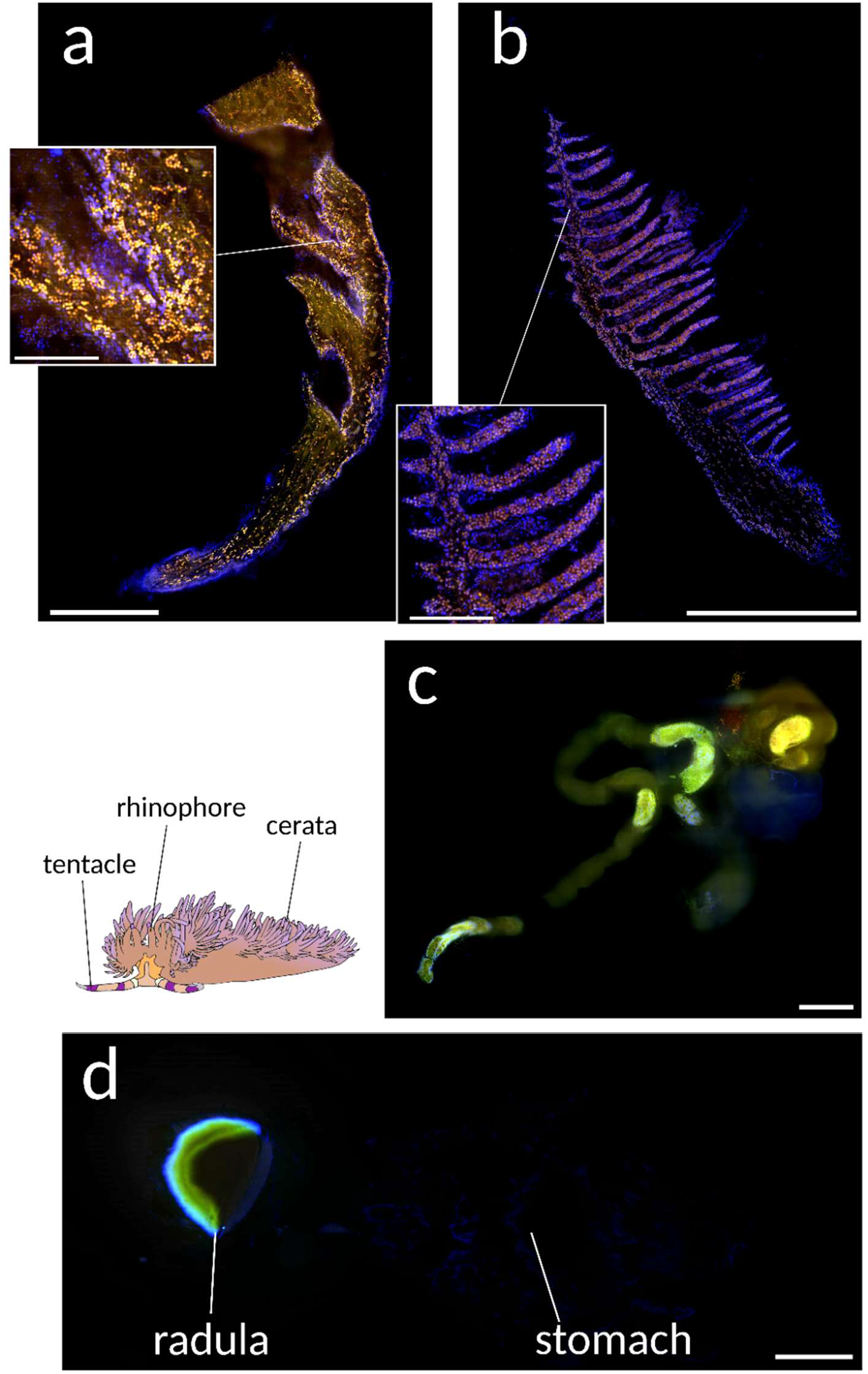
Distribution of Symbiodiniaceae in multiple tissues. a. Tentacle and b. rhinophore, exhibiting abundant algae presence. c. Genital organs and d. radula and stomach, revealing no detectable zooxanthellae. Orange dots represent chlorophyll autofluorescence (Symbiodiniaceae), whereas blue dots indicate cell nuclei (DAPI staining). Scale bar: 1 mm. Scale bar in magnified views of a and b: 250 µm.

### Induction of bleached specimens

We cultured the collected *P. semperi* specimens under fasting conditions to induce the bleaching observed in nudibranchs. A total of 8 individuals, collected in May 2021, exhibited an average survival time of 83 days. Over time, all specimens displayed a progressive paling of body color, ultimately undergoing rapid bleaching and turning completely white 1–2 weeks prior to death (Fig. 2a–c). Concurrent with the bleaching process, a decrease in body length was observed; however, no quantitative analysis of the change in body length was conducted.

**Figure 2.**
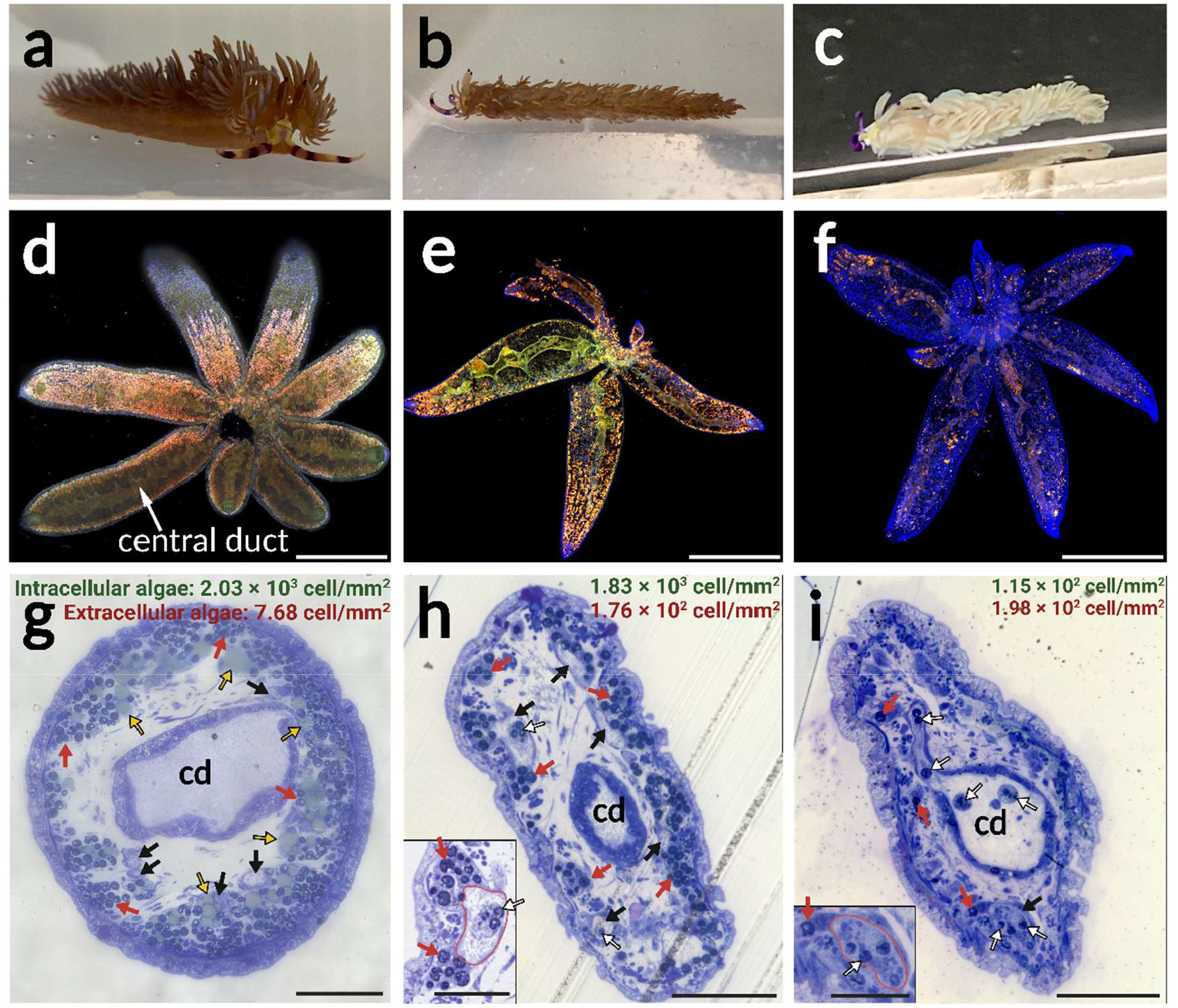
Symbiodiniaceae release through the central duct during body color bleaching *a. P. semperi* immediately after collection, exhibiting a dark-brown body color. b.An unbleached individual following 50 days of starvation, maintaining a brown body color. c.A bleached individual after 91 days of starvation, displaying significantly faded body coloration. d.Fluorescence microscope image of cerata immediately after collection, showing abundant and densely packed Symbiodiniaceae (chlorophyll autofluorescence). Nuclei stained with DAPI in blue. Scale bar: 1 mm. e.Unbleached cerata (50-day fasted) still containing numerous zooxanthellae. Scale bar: 1 mm. f.Cerata from a bleached individual (starved for 50 days) with a significantly reduced density of algae, most of which are located in the central duct. Scale bar: 1 mm. g.Toluidine blue-stained ceras section of an individual immediately after collection. Large amounts of zooxanthellae and lipid droplets are present at the outermost margin and contained within the nudibranch cell. Algae are rarely found in central ducts or fine tubules. The intracellular Symbiodiniaceae density in nudibranch cells (green) and the free extracellular algae density in the lumen of central ducts/fine tubules (red) were calculated for the total area of the ceras section, respectively. Scale bar: 100 µm. h.Ceras section of an unbleached individual (15 days starvation). The lower left shows a magnified view of the fine tubule. In the magnified view, the fine tubule is circled in red. Symbiodiniaceae are abundant around the fine tubules, but some zooxanthellae are released into the lumen of the fine tubules. Lipid droplets are entirely lost. Scale bars: 100 µm (overall view), 50 µm (magnified view). i.Ceras cross-section of a bleached individual (50 days starvation). The lower left displays a magnified view of the fine tubule. Most of the zooxanthellae have been lost, and the remaining ones are found predominantly in the fine tubules and central duct. Scale bar: 100 µm (overall view), 50 µm (magnified view). cd: central duct, black arrow: fine tubules, red arrow: Symbiodiniaceae in nudibranch cells, white arrow: Symbiodiniaceae released into fine tubules and central duct, yellow arrow: lipid droplets.

### Decrease in Symbiodiniaceae density consistent with bleaching of body color

We investigated the changes in algal abundance during body color bleaching by observing chlorophyll autofluorescence in bleached and unbleached nudibranchs using samples of the cerata, a tissue known to retain Symbiodiniaceae and which is easy to collect. The cerata of a 0-day starved (unbleached) individual exhibited a high density of zooxanthellae (Fig. 2d). After 50 days of starvation, the cerata of an unbleached individual still contained numerous algae (Fig. 2e), although their density had decreased. In contrast, the bleached individual displayed a significant reduction in zooxanthellae density (Fig. 2f). At the center of the cerata, a branched digestive gland called the “central duct”^23^ extended toward the outer edge, and Symbiodiniaceae were also observed within its lumen.

### Cross-sectional observation with toluidine-blue staining

We employed optical and electron microscopy to examine the symbiotic structure under different starvation states. Toluidine blue-stained sections revealed that in the cerata, the central duct penetrated the center and branched toward the outer edge in the form of fine tubules.^23^ In the cross-section of ceras from a 0-day fasted individual, Symbiodiniaceae with a density of 2.03 × 10^3^ cells/mm^2^ strongly stained with toluidine blue and numerous lipid droplets were observed within the nudibranch cells (Fig. 2g). In the unbleached individual after 15 days of starvation, the lipid droplets were completely depleted (Fig. 2h), and the intracellular algal density was 1.83 × 10^3^ cells/mm^2^. In the fresh individual, only 1 cell (7.68 cells/mm^2^) of zooxanthellae was released into the lumen of the digestive glands (central duct and fine tubules), whereas 11 cells (1.76 × 10^2^ cells/mm^2^) were released in the 15-day starved individual. Under both 0 and 15-day fasting conditions, intracellular Symbiodiniaceae were present in the intermediate region between the fine tubules and the ceras exothelium. In the bleached individual after 50 days of starvation, intracellular zooxanthellae were significantly reduced to 7 cells (1.15 × 10^2^ cells/mm^2^) (Fig. 2i), and 12 cells (1.98 × 10^2^ cells/mm^2^) were observed in the lumen of the digestive glands. In the tentacle and rhinophore, no central duct-like structures were observed, but fine tubules and algae were present (Supplementary Figs. 1 and 2). In the body, fine tubules and Symbiodiniaceae were distributed just below the surface of the epidermis, with thicker digestive glands observed inside the fine tubules and zooxanthellae. These thick glands were rich in Symbiodiniaceae (Supplementary Fig. 3).

### Cross-sectional observation with TEM

Transmission electron microscopy (TEM) observations provided insights into the ultrastructure of the symbiosis-associated region. In the ceras of the 0-day fasted individual, the epithelial cells of the fine tubules expanded into large clusters, accommodating a substantial number of algae and lipid droplets, thus forming carrier cells^25^ (Fig. 3a). Some Symbiodiniaceae within the carrier cells were observed undergoing cell division (Fig. 3a). In the 15-day starved (unbleached) individual, the carrier cells had lost their lipid droplets and aggregated as the epithelium of the fine tubules (Fig. 3b). Additionally, a small number of zooxanthellae were released alongside their carrier cells into the lumen of the fine tubules (Supplementary Data 1). In the 50-day starved (bleached) individual, the clustered structure of the carrier cells was entirely lost, and the release of algae into the lumen of the central duct and fine tubules had progressed significantly. Some of the ejected Symbiodiniaceae were enveloped in a form of carrier-cell debris (Fig. 3c). A few Symbiodiniaceae still remained within the epithelium of the fine tubules, some of which had collapsed (Fig. 3d). The rhinophore and the tentacle were not penetrated by the central duct but were infiltrated only by the fine tubules.

**Figure 3.**
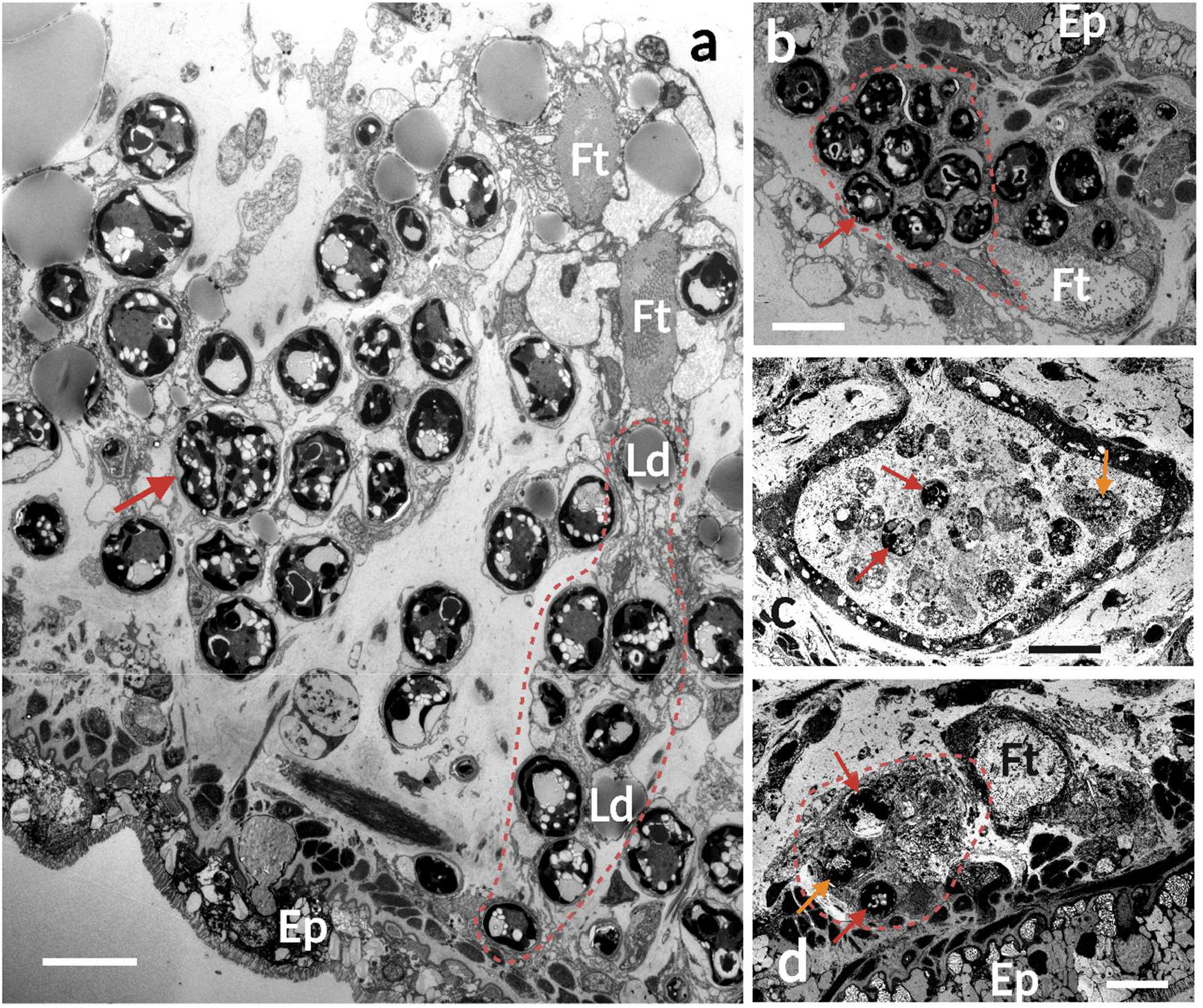
TEM images of nudibranch cerata under each starvation condition; refer to Supplementary Data 1 and 2 for complete images of b–d. a.Longitudinal section of a freshly collected individual, exhibiting a clustered structure with the epithelium of the fine tubules broadly expanded as a carrier cell. Numerous Symbiodiniaceae and lipid droplets are stored inside. The algae indicated by the arrow are undergoing cell division. Scale bar: 10 µm. b.Cross-section of a 15-day fasted (unbleached) individual, displaying the loss of lipid droplets from the carrier cell and shrinkage of its clustered structure. Aggregation of zooxanthellae around the fine tubules is observed. Scale bar: 10 µm. c.Central duct in the cross-section of a 50-day fasted (bleached) individual’s ceras, revealing the release of algae accompanied by cellular debris. The Symbiodiniaceae indicated by the orange arrow are wrapped in a carrier cell-like structure, and the algal cell has collapsed. Scale bar: 20 µm. d.Cross-section of a 50-day fasted (bleached) individual’s ceras, demonstrating the complete loss of the clustered structure of the carrier cells and a substantial reduction in the density of internal algae. The Symbiodiniaceae indicated by the orange arrow exhibit a collapsed cellular structure. Scale bar: 10 µm. Ft: fine tubules, Ld: lipid droplet, Ep: ceras epidermis, dashed line: carrier cell.

### Observation of symbiosis with SEM

The 3D structure of the cerata was investigated through SEM observation of a split section of the cerata. Consistent with previous observations, a central duct runs through the center, accompanied by fine tubules (Fig. 4a). The structure of these 2 types of digestive gland-derived ducts differed: the central duct exhibited a mesh-like pattern, whereas the fine tubules were hollow. Three-dimensional clusters of carrier cells containing Symbiodiniaceae and lipid droplets were observed external to the fine tubules in the 0-day starved individual (Fig. 4b, c). Within the lumen of the fine tubule, a collection of algae was observed, partially engulfed by the epithelial cells (Fig. 4d). In the unbleached individual subjected to 15 days of fasting, algal cells remained present in large numbers at the outer margin of the ceras but were also detected as having been released into the lumen of the central duct (Fig. 4e).

**Figure 4.**
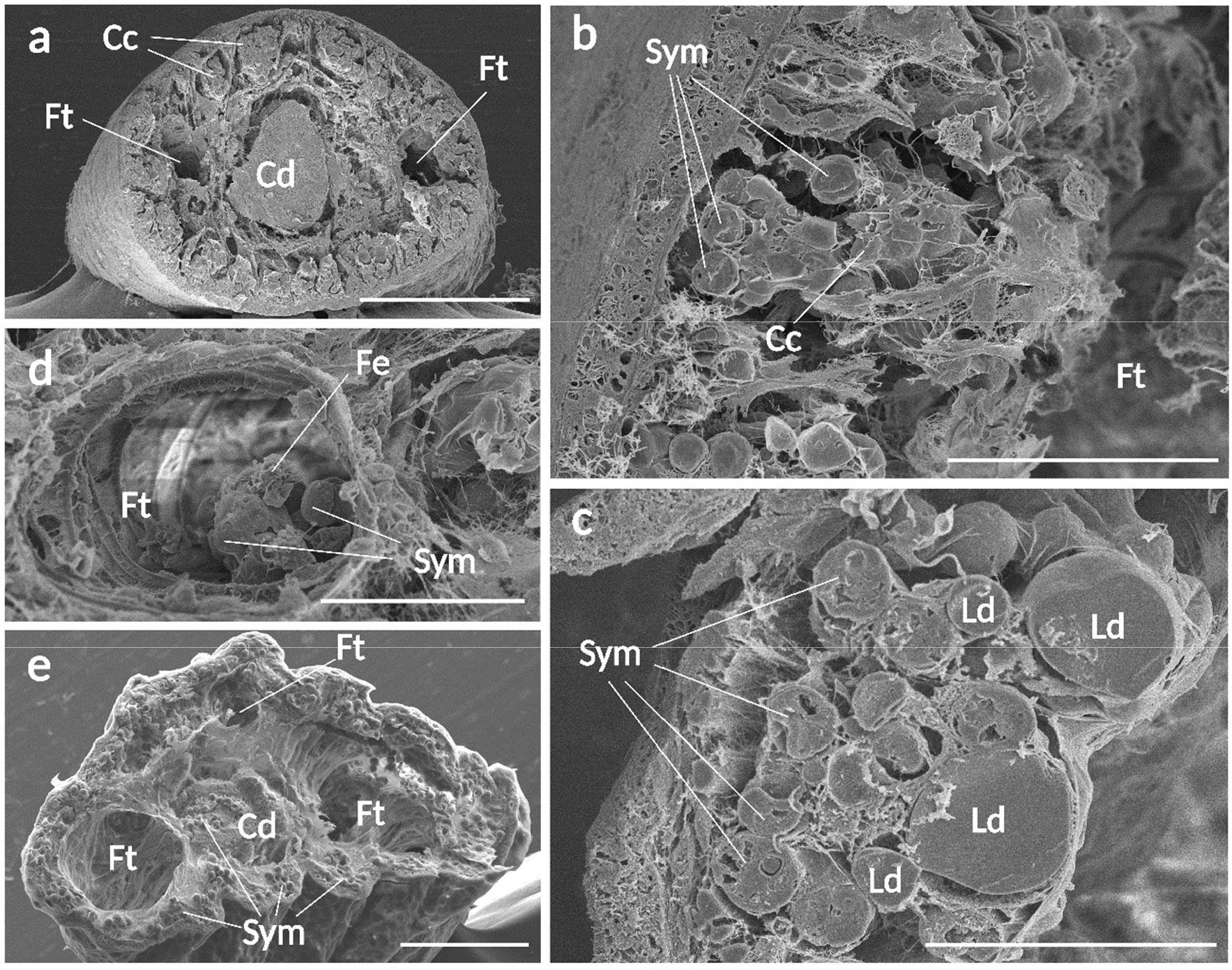
SEM images of transversely sectioned cerata. a–d: 0-day fasted individual, e: 15-day fasted individual (unbleached). a.Overview of a ceras. The central duct runs through the center, followed by fine tubules. Carrier cells are located at the outermost margin. Scale bar: 200 µm. b.Cluster-like arrangement of carrier cells surrounding fine tubules. Scale bar: 50 µm. c.Cross-section of a carrier cell. Multiple lipid droplets and Symbiodiniaceae are visible. Organelle structures can also be detected in the zooxanthellae. Scale bar: 30 µm. d.A collection of algae within a fine tubule, appearing to be engulfed by epithelial cells. Scale bar: 40 µm. e.Overview of a ceras in a 15-day fasted (unbleached) specimen; the carrier cells have shrunk, and the entire ceras appears atrophied. Symbiodiniaceae can be observed in the central duct as well as at the outer margin of the ceras. Scale bar: 100 µm. Cd: central duct, Ft: fine tubule, Fe: epithelium of fine tubules, Cc: carrier cell, Sym: Symbiodiniaceae, Ld: lipid droplet.

## Discussion

### Highly developed symbiotic structure in *P. semperi*

Our thorough microscopic investigation has illuminated the intricate histological structure of the advanced symbiotic system present in *P. semperi*. The fine tubules,^23^ which are characteristic of highly branched digestive glands, consistently permeate just below the superficial body tissues, whereas no analogous structures can be observed within the internal organs of the organism. This suggests that *P. semperi* has refined its algal symbiosis system to be primarily situated on the body surface, where light conditions are most favorable for the photosynthetic processes of the zooxanthellae.

In freshly collected healthy specimens, the epithelial cells of the fine tubules were observed to expand, forming a clustered structure housing multiple algal cells and lipid droplets. This sophisticated structure is likely the “carrier” cell described by Kempf.^25^ Although the carrier cells appeared unattached in cross-sectional views, as previously reported by Kempf (Supplementary Data 1 and 2), longitudinal sections revealed that the majority of these cells maintained connections to the fine tubules (Fig. 3a, Supplementary Data 3). This suggests that the carrier cells are not entirely independent from the fine tubules; rather, they exhibit an elongated and distended morphology. Within the carrier cells, lipid droplets tended to be situated closer to the fine tubules in the inner part of the body, in comparison to the distribution of Symbiodiniaceae (Fig. 2g). The clustered structure of the carrier cells and the strategic arrangement of lipid droplets within these cells help position the zooxanthellae at the outermost edge of the body, providing a favorable light environment for the algae. The observation that some Symbiodiniaceae were dividing (Fig. 3a) further supports the notion that carrier cells offer a beneficial niche for the algae. The tufted structure may additionally assist the sea slugs in accommodating a larger number of zooxanthellae. In the genus *Berghia*, most commonly utilized in nudibranch-algal symbiosis studies, which is considered a primitive species with respect to algal symbiosis,^21^ the digestive gland epithelial cells merely hold the zooxanthellae and do not swell.^26^ In contrast, the symbiotic morphology of *P. semperi* is deemed more sophisticated in terms of Symbiodiniaceae harboring capacity and light conditions. This supports the prevailing theory that this genus represents one of the nudibranchs with the most highly developed symbiotic capabilities.^23^

The sophisticated partnership between Symbiodiniaceae and their hosts involves the transfer of photosynthetic products, often in the form of lipid droplets. In coral and foraminiferal hosts, these lipid droplets are believed to be the primary photosynthetic products of the algae.^27–29^ Previous research^22^ suggests that lipids and glycerol constitute the main components of photosynthates translocated from Symbiodiniaceae to *Pteraeolidia*, supporting the notion that the observed lipid droplets in carrier cells originate from zooxanthellae. Intriguingly, the lipid droplets in *P. semperi* were significantly larger than those documented in other hosts, with diameters reaching up to 30 µm (Fig. 2g), as opposed to the 1–5 µm typically observed in corals^27,28^ and foraminifera.^29^ Conversely, in the sacoglossan *Elysia chlorotica*, which possesses “kleptoplasty” capabilities, chloroplasts generate lipid droplets within the sea slug’s body that closely resemble those of *P. semperi* in both morphology and size.^30^ This suggests that the presence of these sizable lipid droplets may be a unique characteristic of photosymbiosis in Gastropoda. Sea slug hosts, possessing greater mobility than other host species, are likely to acquire a substantial amount of heterotrophic nutrients. Whereas corals are thought to rely on the photoautotrophic nutrition provided by Symbiodiniaceae for 65–85%^31,32^ of their metabolic requirements, *P. semperi* and *E. chlorotica* may be less dependent on their symbiotic partners, storing a significant proportion of the photosynthates they receive from zooxanthellae within their carrier cells rather than utilizing them directly. To further elucidate the processes of lipid droplet formation and metabolism within this symbiotic relationship, as well as the specific roles these droplets play, additional investigations involving compositional analysis, radioisotope tracing, and analysis with NanoSIMS are warranted.

### The bleaching process in *P. semperi*

The induction of body color bleaching in *P. semperi* was achieved through a period of starvation. Although this study did not undertake a quantitative analysis of the shifts in body color, future investigations should address this aspect, in conjunction with other influential parameters such as water temperature and illumination. A comparison of algal abundance before and after bleaching revealed that, akin to corals, bleaching in *P. semperi* results from the loss of symbiotic zooxanthellae. Furthermore, by examining the ultrastructure under 3 distinct fasting conditions, we were able to deduce the cellular mechanisms involved in each stage of the *P. semperi–*Symbiodiniaceae symbiosis (Fig. 5).

**Figure 5.**
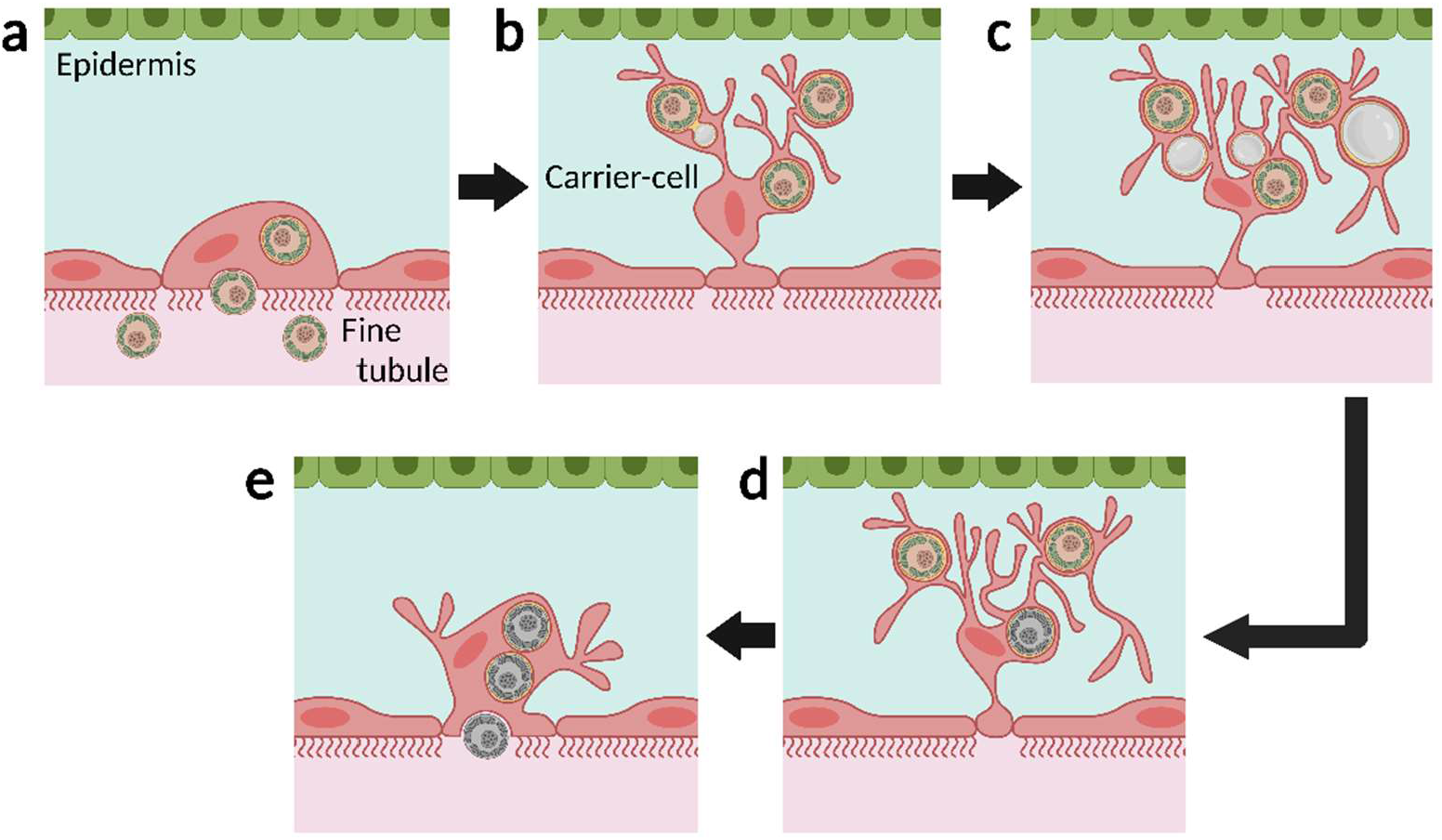
Proposed cellular mechanism underlying the *P. semperi–*Symbiodiniaceae symbiosis a.Symbiodiniaceae are endocytically engulfed by the epithelial cells of the fine tubules. b.Epithelial cells of the fine tubules undergo significant expansion toward the epidermis, forming carrier cells. c.Algae, situated just beneath the body surface and held by carrier cells, perform photosynthesis, leading to the formation of lipid droplets within the carrier cells. d.Upon the onset of starvation, the sea slug host metabolizes the lipid droplets to supplement its energy requirements. e.As the symbiosis deteriorates and zooxanthellae perish, the carrier cells contract and release the zooxanthellae into the lumen of the fine tubules.

### Acquisition of Symbiodiniaceae

Kempf^25^ described a “dark brown organ” located in an extension of the digestive gland and free “carrier” cells situated in the intermediate region between the digestive gland and the cerata epithelium in *Pteraeolidia*. He postulated that algae within the lumen of the digestive gland could be conveyed into the extracellular matrix via the dark brown organ, ultimately transforming into carrier cells. Contrarily, our investigation did not uncover any organ analogous to the dark-brown organ. Moreover, during SEM analysis (Fig. 4d), we observed that a fraction of the zooxanthellae population was integrated into the epithelium of the fine tubules. Although it is unclear whether this signifies a moment of acquisition or release of Symbiodiniaceae, these findings imply that the epithelial cells of the fine tubules engage in direct algal exchange. Given that carrier cells are likely fine tubules’ epithelial cells, and zooxanthellae are presumably engulfed by the digestive gland epithelium via phagocytosis in primitive symbiotic nudibranchs, it seems plausible that there is no intermediary organ and that the zooxanthellae within the lumen of the fine tubules are directly assimilated into the carrier cells. The genus *Pteraeolidia* has recently been divided into 2 species, *P. semperi* and *P. ianthina*, and the potential existence of cryptic species is still being considered.^33^ It cannot be discounted that the dark brown organ may serve a functional role in the specimens employed by Kempf (collection site unspecified, but potentially from Hawaii). Nonetheless, in *P. semperi* at least, orally introduced Symbiodiniaceae are dispersed throughout the organism via the highly branched digestive gland, where they are phagocytically captured by epithelial cells in the fine tubules, which extend from the gland (Fig. 5a).

### Establishment of symbiosis

As the Symbiodiniaceae algae are assimilated, the fine tubules’ epithelial cells undergo a tufted expansion toward the body surface, evolving into carrier cells and maintaining the algae proximate to the tissue’s outer boundary (Fig. 5b). The algae, residing within the carrier cells, perform photosynthesis and subsequently discharge photosynthetic byproducts as lipids into the carrier cells, giving rise to lipid droplets (Fig. 5c). It is conjectured that during periods of starvation, the sea slug ingests these lipid droplets, harnessing them for metabolic sustenance.

### Dissolution of symbiosis

During prolonged periods of starvation, lipid droplets become depleted (Fig. 5d), leading to the conjecture that carrier cells revert to their original state as digestive gland epithelial cells and release the algae into the gland lumen (Fig. 5e). The process of release may encompass not only exocytosis but also instances in which the carrier cells entirely detach into the lumen (Fig. 3c). Before and after release, a substantial proportion of zooxanthellae were found to have expired, exhibiting disintegrated cellular structures. The precise factors that trigger the release of Symbiodiniaceae remain a subject for further discussion, such as whether the famished sea slug actively digests the algae or if it simply excretes algae that have deteriorated or succumbed to natural death due to aging.

Throughout the elucidation of these mechanisms, it becomes unequivocally clear that the epithelial cells of the fine tubules, namely carrier cells, play an indispensable role in the symbiotic relationship. Despite their unique morphology, the carrier cells demonstrate functional similarities to the gastrodermal cells of corals, particularly regarding the intracellular uptake and extracellular release of Symbiodiniaceae, as well as the formation of lipid droplets by algae. This correlation reinforces the relevance of the *P. semperi* model in investigations pertaining to coral–Symbiodiniaceae symbiotic mechanisms. In the context of coral-algal symbiosis, the participation of pattern recognition receptors and a variety of ion transporters is documented at the symbiotic interface,^12^ suggesting that analogous molecular mechanisms may be present in carrier cells. Although coral bleaching is typically triggered by thermal or light stress,^34^ this study observed bleaching in the sea slug due to starvation. Exploring the molecular mechanisms within symbiotic cells under distinct stressors could yield valuable insights. Nevertheless, the current state of knowledge in molecular biology concerning the *P. semperi–*algal symbiotic system is virtually nonexistent and remains incomparable to that of coral symbiotic systems. Future investigations into molecular mechanisms will necessitate the application and evaluation of diverse molecular biological approaches, focusing on carrier cells under a range of conditions.

## Conclusion

The symbiosis between Symbiodiniaceae algae and sea slugs has been acknowledged for an extended period, yet a comprehensive understanding of the associated structures and mechanisms has remained elusive. In this research, we have rigorously delineated the intricately developed landscape of the *P. semperi–*Symbiodiniaceae symbiotic relationship. Moreover, through examining the process of zooxanthellae loss (bleaching), we have inferred the underlying histological mechanisms. This information serves as a cornerstone for molecular investigations into sea slug–algal symbiosis, encompassing transcriptomics and metabolomics, and it is expected to contribute to the overarching endeavor of deciphering the complex mechanisms governing coral–Symbiodiniaceae symbiosis.

## Acknowledgments

We extend our gratitude to Dr. Yusuke Kijima for his valuable advice on improving the manuscript. We also express our appreciation to the Technology Advancement Center at the Graduate School of Agricultural and Life Sciences at the University of Tokyo for providing access to electron microscopy facilities. This research was conducted with the support of the JSPS Grant-in-Aid for Scientific Research (A) [20H00429] and the Sasakawa Scientific Research Grant [2022-4042]. The figures included in this paper were created using BioRender (BioRender.com).

## Author Contributions

Hideaki Mizobata conceived the research concept. Hideaki Mizobata, Kentaro Hayashi, and Ryo Yonezawa undertook the collection and cultivation of *P. semperi* specimens. Hideaki Mizobata was responsible for overseeing the whole-mount observation process. Kenji Tomita and Hideaki Mizobata carried out the electron microscopy analyses. All authors engaged in thorough discussions regarding the interpretation of the results. Hideaki Mizobata composed the manuscript, and all authors contributed to its revision and granted their approval for the publication of the final version.

## Supplementary Figures

1.Cross-section of a tentacle from an unbleached individual subjected to 15 days of starvation, stained with toluidine blue.

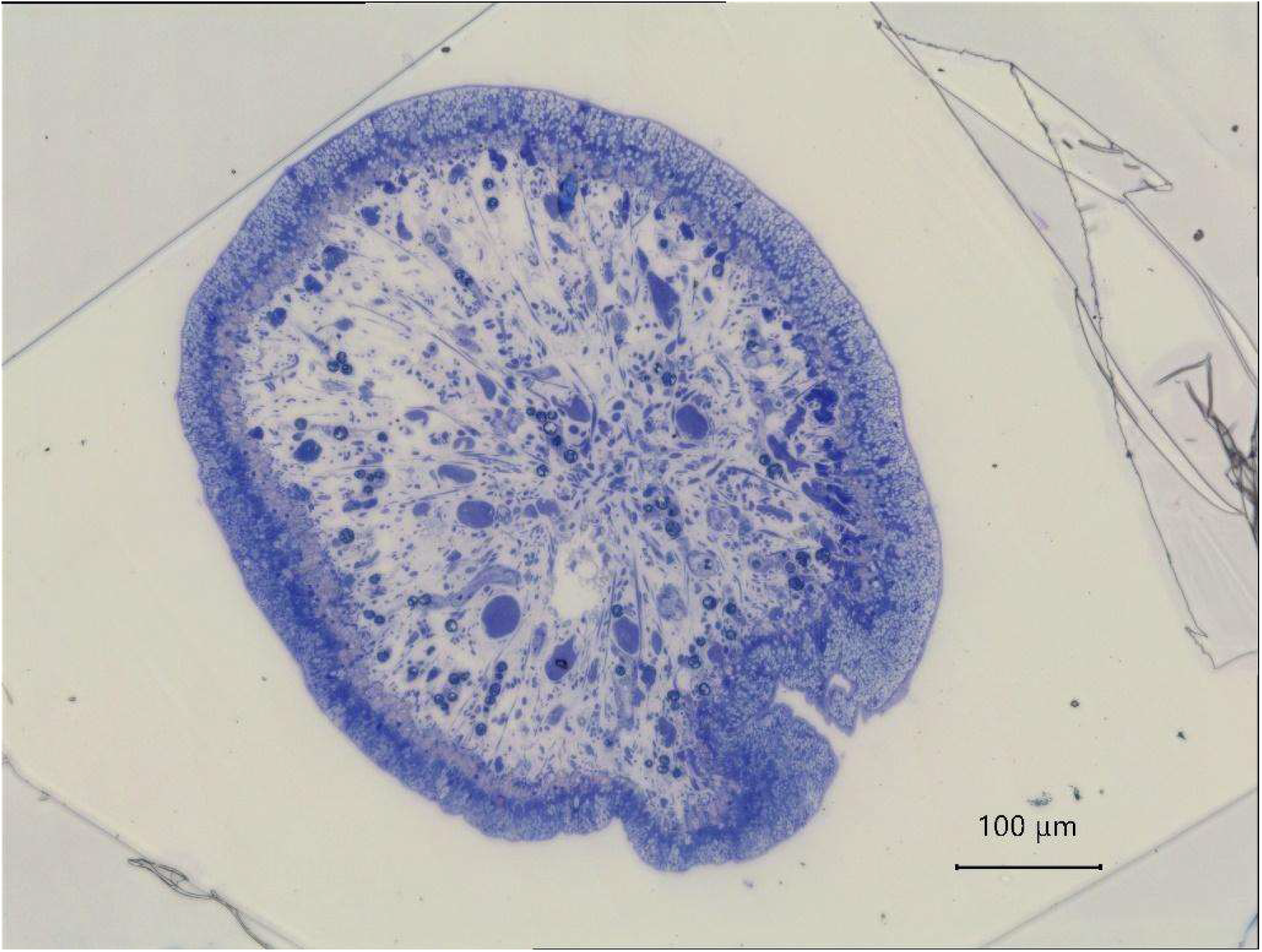

2.Cross-section of a rhinophore from an unbleached individual subjected to 15 days of starvation, stained with toluidine blue.

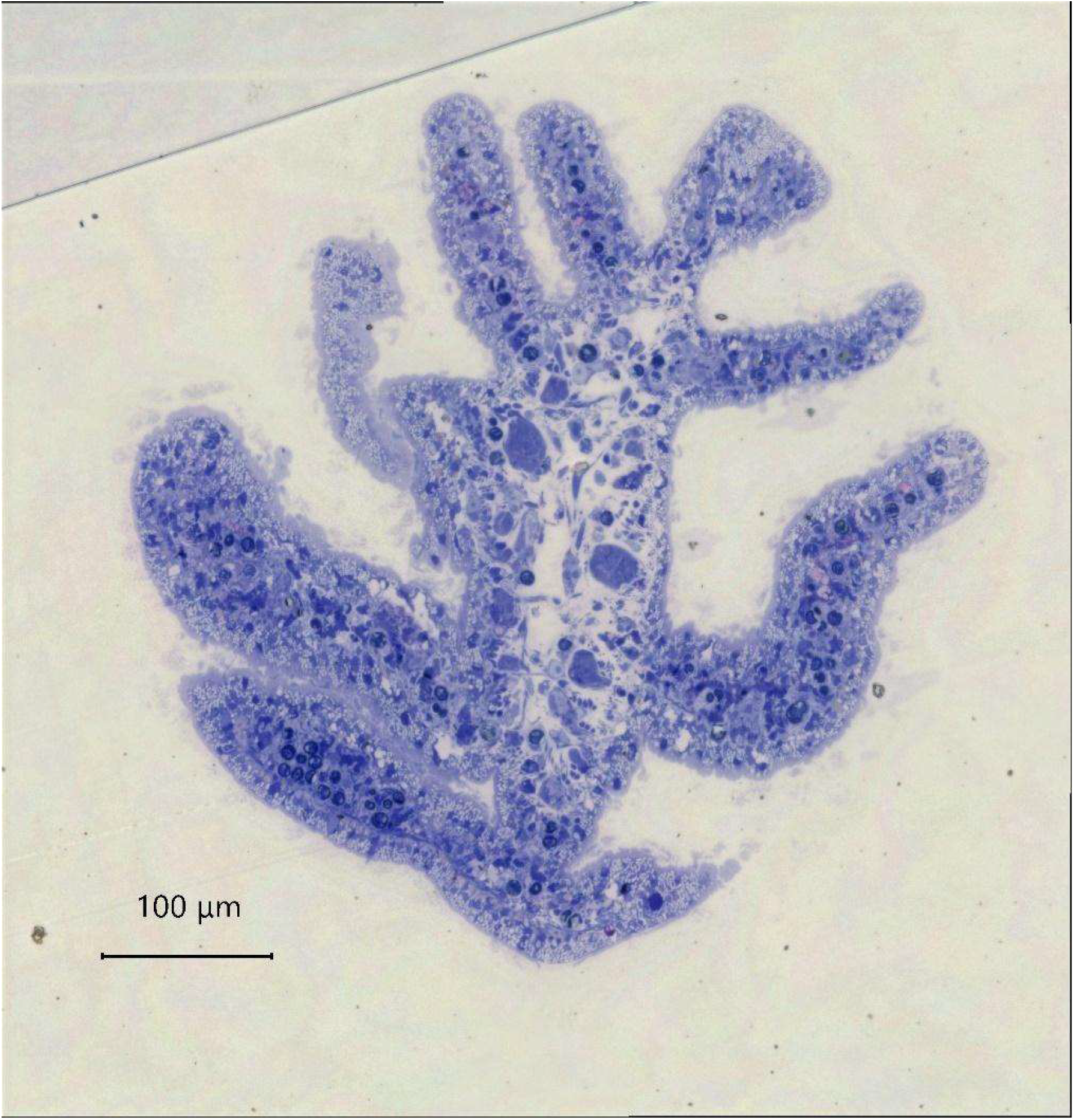

3.Cross-section of a body from an unbleached individual subjected to 15 days of starvation, stained with toluidine blue.

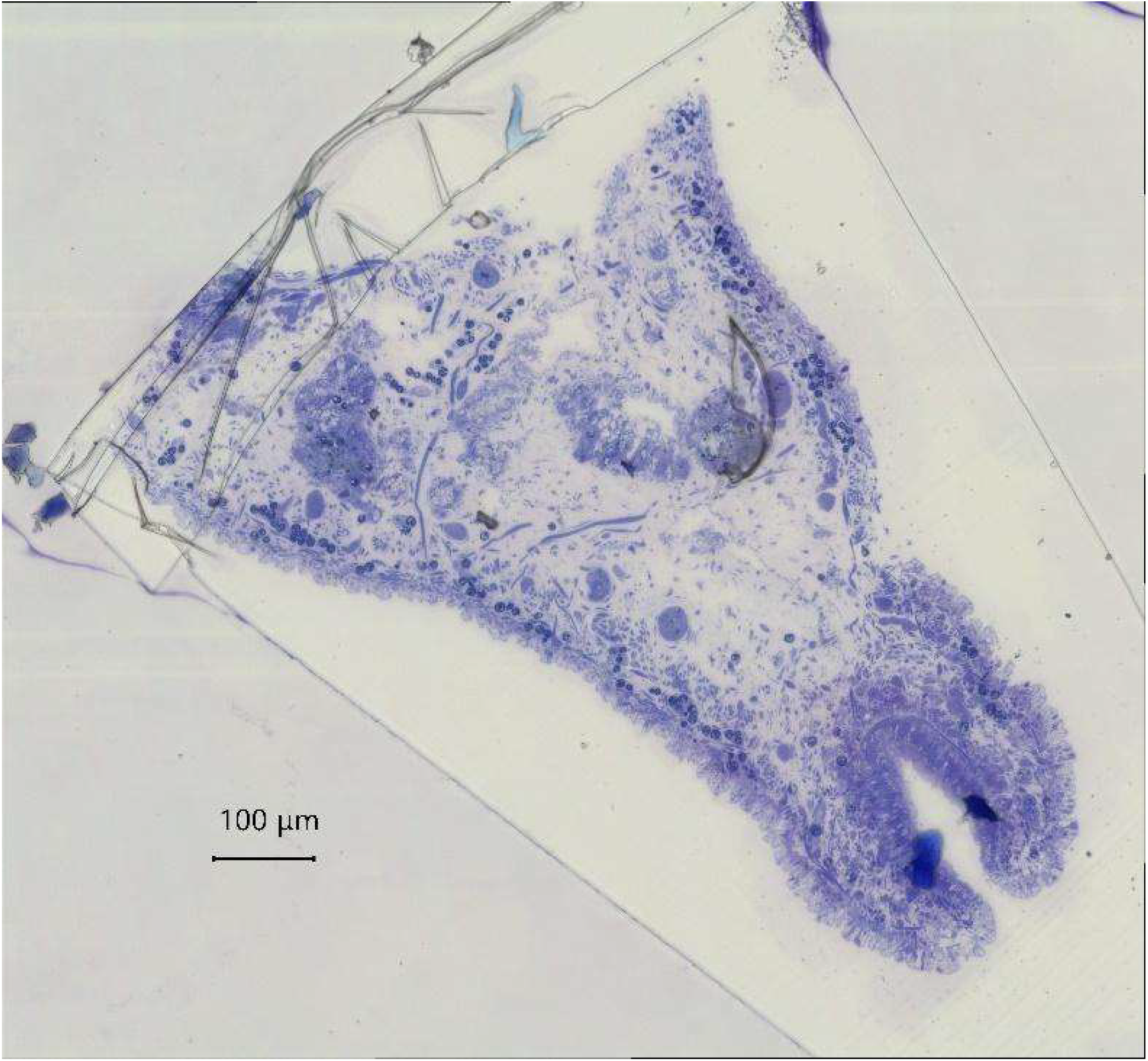

## Supplementary Data

1.Combined TEM images of a cerata cross-section from an unbleached individual subjected to 15 days of starvation.

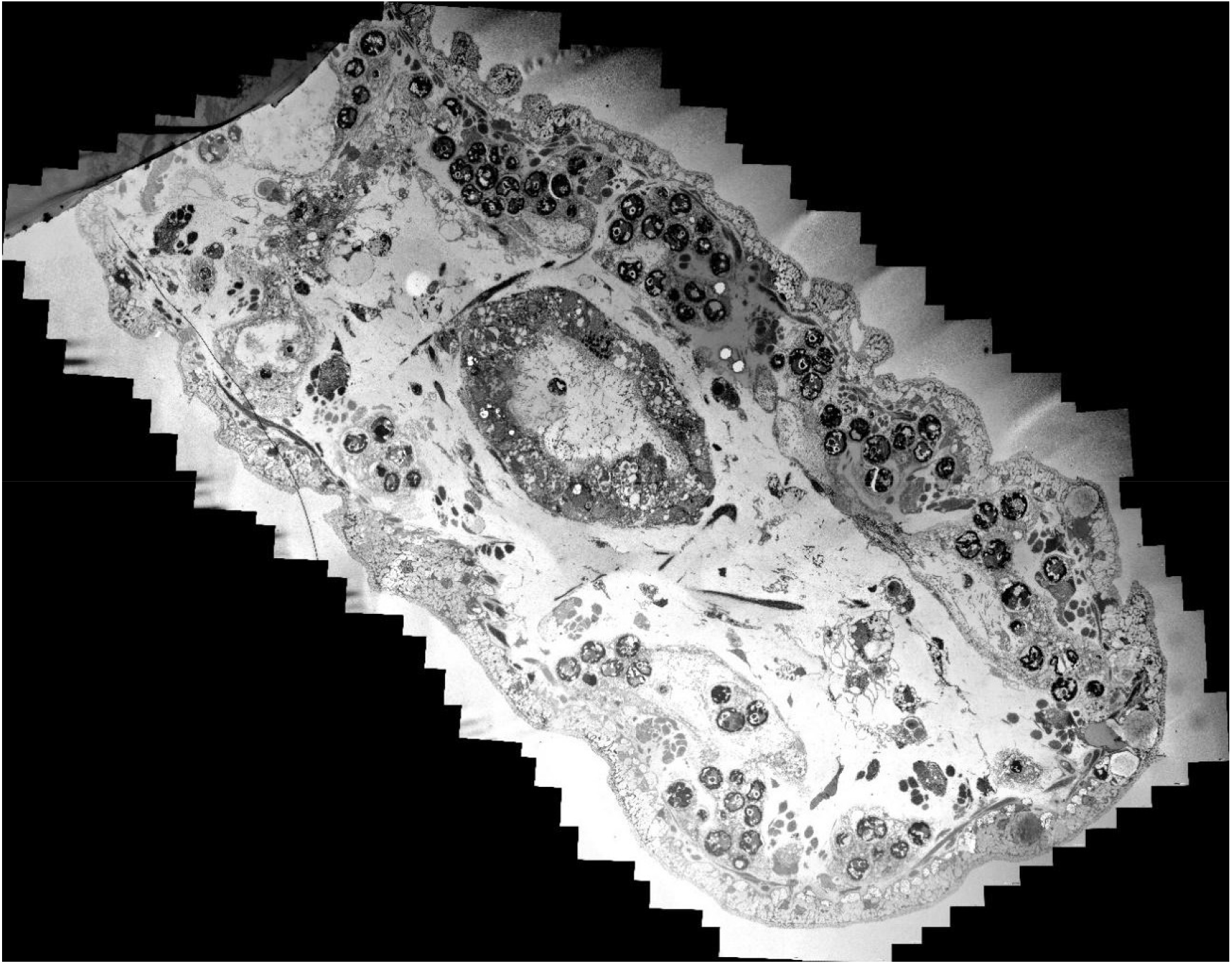

2.Combined TEM images of a cerata cross-section from a bleached individual subjected to 50 days of starvation.

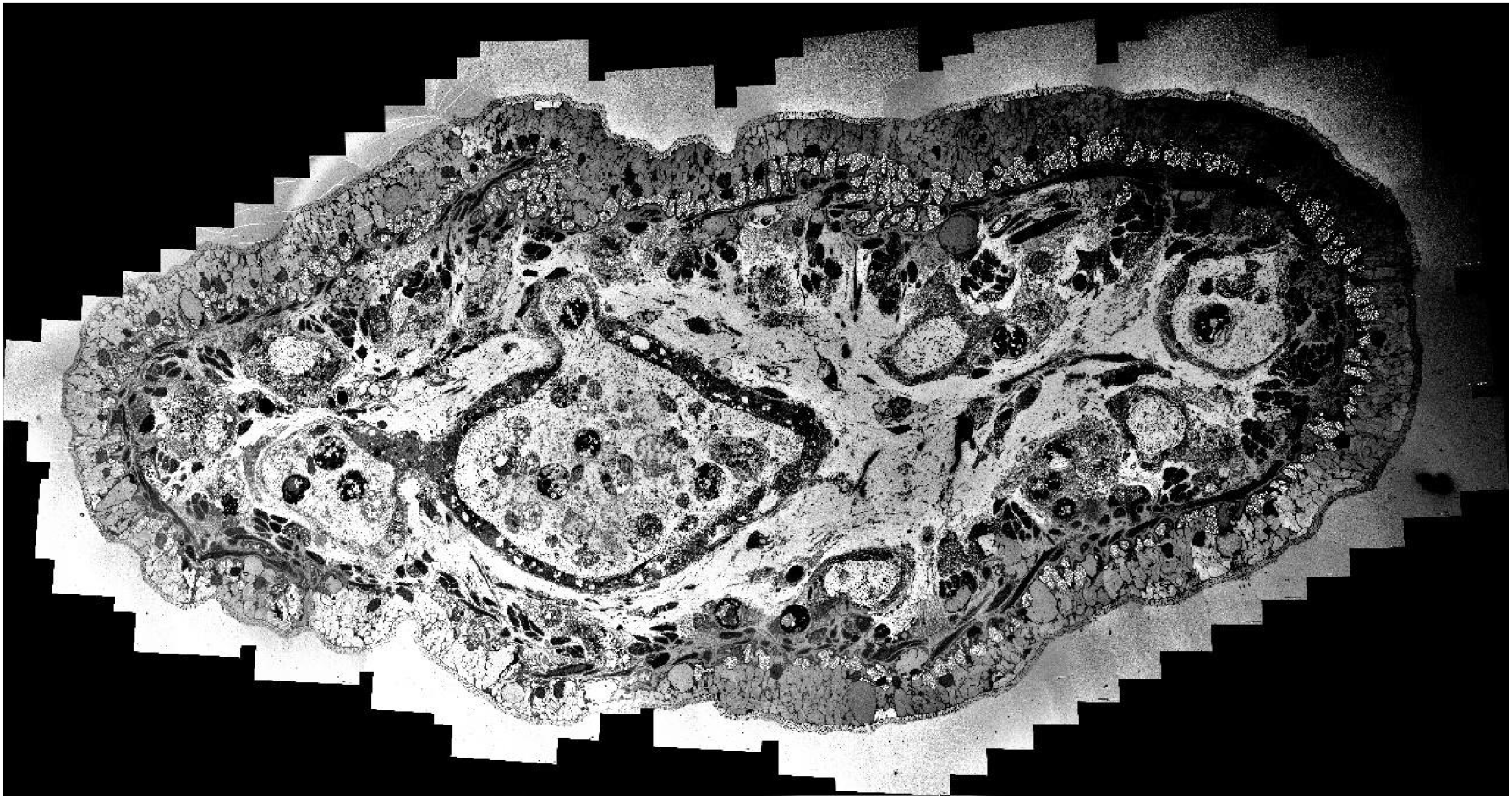

3.Combined TEM images of a cerata longitudinal section from a fresh individual (unbleached).

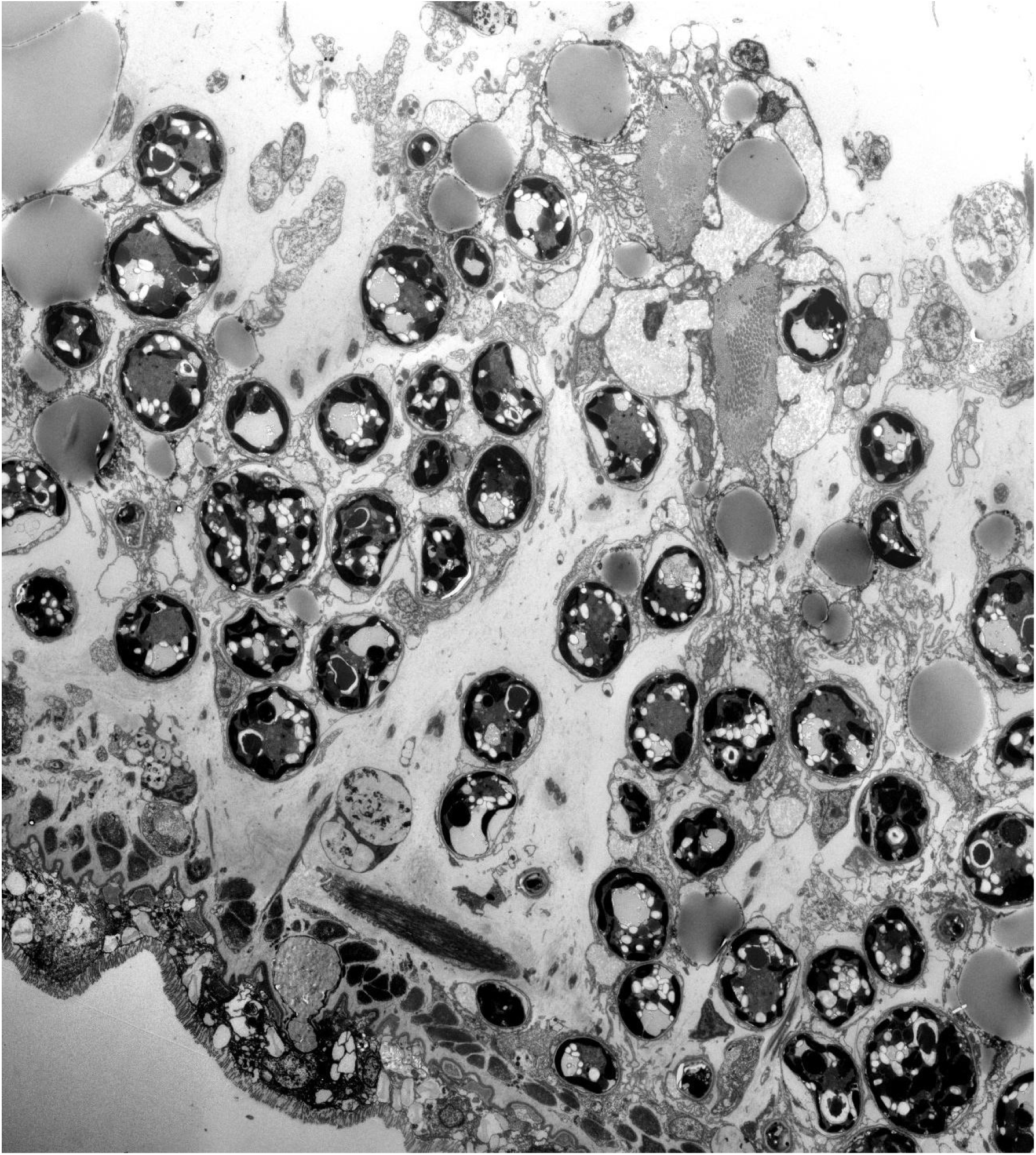

